# Joint Genotype- and Ancestry-based Genome-wide Association Studies in Admixed Populations

**DOI:** 10.1101/062554

**Authors:** Piotr Szulc, Malgorzata Bogdan, Florian Frommlet, Hua Tang

**Affiliations:** Faculty of Mathematics, Wroclaw University of Technology, Poland; Institute of Mathematics, Wroclaw University, Poland; Department of Medical Statistics, CEMSIIS, Medical University of Vienna; Departments of Genetics and Statistics, Stanford University

**Keywords:** Admixture Mapping, Model Selection, Multiple Regression, Quantitative Trait

## Abstract

In Genome-Wide Association Studies (GWAS) genetic loci that influence complex traits are localized by inspecting associations between genotypes of genetic markers and the values of the trait of interest. On the other hand Admixture Mapping, which is performed in case of populations consisting of a recent mix of two ancestral groups, relies on the ancestry information at each locus (locus-specific ancestry).Recently it has been proposed to jointly model genotype and locus-specific ancestry within the framework of single marker tests. Here we extend this approach for population-based GWAS in the direction of multi marker models. A modified version of the Bayesian Information Criterion is developed for building a multi-locus model, which accounts for the differential correlation structure due to linkage disequilibrium and admixture linkage disequilibrium. Simulation studies and a real data example illustrate the advantages of this new approach compared to single-marker analysis and modern model selection strategies based on separately analyzing genotype and ancestry data, as well as to single-marker analysis combining genotypic and ancestry information. Depending on the signal strength our procedure automatically chooses whether genotypic or locus-specific ancestry markers are added to the model. This results in a good compromise between the power to detect causal mutations and the precision of their localization. The proposed method has been implemented in R and is available at http://www.math.uni.wroc.pl/~mbogdan/admixtures/.

## 1 Introduction

Genome-wide association studies have proven to be a powerful approach for mapping loci that underly complex traits. GWAS data are most commonly analyzed by testing each marker individually. Given the large number of markers involved it is crucial to carefully address the problem of multiple testing. Classical approaches include the Bonferroni correction, which controls the Family Wise Error Rate (FWER), or the Benjamini-Hochberg correction ([4], BH), aimed at controlling the False Discovery Rate (FDR). In these methods p-values for single-marker tests are adjusted, taking into account the total number of tests.

When a trait is polygenic, the power of single-marker testing can be improved by adjusting for variation at known trait loci. This partially explains the improved power of the popular mixed effects approaches, such as EMMAX [27], which model polygenic background as random effects. Alternatively one can build multi-locus models, which have the additional advantage of opening the possibility to incorporate gene-gene or gene-environment interaction terms. A fairly large number of model based algorithms for complex trait mapping are now available (see for example [23, 14] and references given there), but not all of them treat the resulting model selection problem rigorously. Regularization based methods, such as LASSO, focus on prediction, and post-selection statistical inference with respect to each selected variable is still an open question. In contrast the methods developed in [19] and [14] for GWAS are based on modifications of the Bayesian information criterion called mBIC and mBIC2. These were specifically designed to control FWER and FDR, respectively, in a high dimensional setting. Here we want to adapt specifically the FDR controlling criterion mBIC2 to the problem of admixture mapping.

Mapping complex traits in populations that have experienced recent admixture, such as the African American and Hispanic populations, are particularly challenging for two reasons. First, recent genetic admixture creates linkage disequilibrium (LD) between unlinked loci, giving rise to spurious association [9, 10, 20]. In GWAS, various methods have been developed, which use high-density genotype data to infer individual-level ancestry proportions; adjusting these ancestry proportions offers an effective solution to eliminate confounding due to population stratification [30, 44]. A second challenge, which has no satisfactory solution, is that minority cohorts are much smaller than the available European cohorts. Under a polygenic genetic architecture there is a large number of loci each of which contributes some moderate effects; as a result, for a trait variant with comparable allele frequency and allelic size across populations, the statistical power of detecting this variant is much lower among African Americans or Hispanics than in the European population.

Improving the power of mapping complex traits in recently admixed minority populations is an important goal for several reasons. First, admixed individuals derive their genome from multiple ancestral populations, and therefore offer opportunities to uncover trait variants that are not polymorphic in a single population. Second, even when a trait variant is shared across populations, its allelic effect may vary. Therefore, for genetic risk assessment, it is important to characterize the effect of each trait variant in the relevant population. Third, admixed populations enable mapping of trait loci that underlie population-level trait difference. Admixture mapping, which seeks genomic regions where the phenotype is statistically associated with the ancestry origin of the chromosomal segment, is particularly powerful to map trait variants with disparate allele frequencies in the ancestral populations [24, 48, 51]. Using high-density genotype data, the ancestry information at each locus along the different chromosomes, referred to as local ancestry, can be accurately inferred using a number of computational approaches [33, 35, 42, 44]; in this study, we assume that local ancestry is known without error.

It was first proposed in [46] to combine genotype-based and ancestry-based tests. There it was shown that the two tests provide complementary information and that adding admixture mapping to the genotype-based tests does not significantly increase the burden of multiple-testing. However, this approach was developed for the parent-trio TDT test and cannot be easily generalized for GWAS based on unrelated individuals. Subsequently, Shriner et al [37] proposed a different testing procedure that combines genotype and ancestry information, using an adhoc “Bayesian” approach which is rather difficult to justify. The method uses the posterior probabilities from admixture analysis as the priors for genotype based tests and relies on the specific selection of the prior distributions on the number of causal mutations and the magnitude of their effects. Our simulation results show that this way of combining ancestry and genotype information can only improve the power of genotype and admixture mapping in situations where both approaches are capable of detecting the signal, otherwise it performs rather badly.

In this study we develop a rigorous procedure for building multi-locus models for complex traits in population-based GWAS, which combines genotype and ancestry information in admixed populations. To this end we consider regression models with two sets of candidate explanatory variables: one set (*X*) representing the genotype of each SNP and a second set (*Z*) representing the local ancestry at the location of the genotyped SNPs. When building models it is important to be aware of the extremely different correlation structure between *X* variables and between *Z* variables, respectively. Correlation between neighboring genotype markers is governed by linkage disequilibrium (LD), which generally decays rapidly due to historic recombination. In contrast, correlation between local ancestry depends on recombination *after admixing;* in recently admixed populations, such as African Americans or Hispanics, correlation in local ancestry decays much more slowly compared to LD between genotypes.

We introduce an FDR-controlling modification of BIC which properly accounts for the differential correlation structure by introducing separate penalty terms for the ancestry and genotype variables. Specifically we will make use of an ‘effective number’ of ancestry state variables to specify the penalty for the *Z* variables. After formally introducing the new selection criterion we compare its performance with the following competing procedures: Single-marker tests, multi-locus models using only genotype information or only local ancestry, respectively, and the BMIX procedure [37]. To this end we perform a comprehensive simulation study under complex genetic models and also reanalyze GWAS data of HDL cholesterol in an African American cohort.

## 2 Methods

### 2.1 Genotype and Admixture mapping

Our goal is the identification of DNA regions harboring “causal” mutations based on a sample of n unrelated individuals from the admixture of two distinct populations. We will focus on GWAS with quantitative traits, where the measurement of the trait for the i-th individual is denoted as *yi, i* ∈ {1,…, *n*}. Furthermore, we assume that for each individual the genotypes of *p* SNPs as well as the corresponding locus specific ancestry are known. The genotype for the *j*th SNP of the *i*th individual is coded as

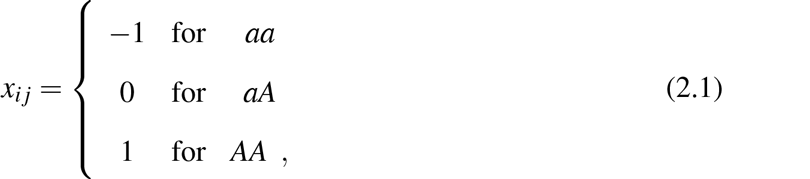

where *a* and *A* generically denote the two variants of each SNP. For our purposes we do not have to know which one is the wild type and which one the mutation. Similarly, the ancestry status of the *i*th individual at the *j*th SNP location is coded as

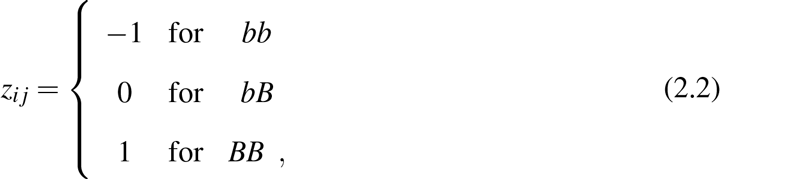

where *b* and *B* refer to the two different ancestral populations. We will use the notation *X_j_* and *Z_j_* for the underlying random variables of genotype and ancestry state at location *j*.

The simplest way to perform GWAS is to test each genotype or ancestry marker individually for association with the trait in question. To eliminate spurious associations due to population stratification, the genome-wide ancestry, 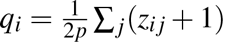, is included in the model as a covariate [34]. Thus, the standard single-marker genotype-phenotype association test uses the model

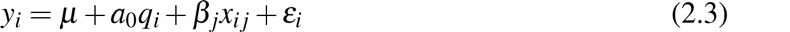

to test the hypotheses *H_xj_*: *β_j_* = 0, for *j* = 1,…, *p*. Likewise, admixture mapping uses the model

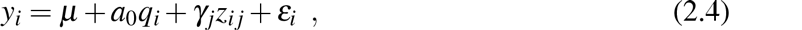

to test the hypotheses *H_zj_*: *γ_j_* = 0. We will use for all models the generic notation *ε_i_* for error terms and will always assume that they are independent and normally distributed, *ε_i_* ~ N(0, σ^2^).

Multi-locus models including only genotypic effects can be written as extensions of (2.3) in the form

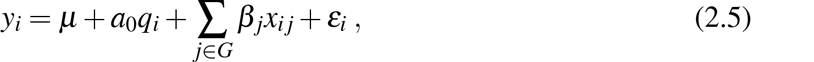

where *G* specifies the model by indexing the subset of markers which might influence the trait. We will also write *X_G_* for the submatrix of *X* which includes only the columns corresponding to the index set *G*. Important SNPs can then be localized by looking for that model which minimizes some model selection criterion which balances the complexity of the model and its fit to the data. One of the most popular tools for this task is the Bayesian Information Criterion (BIC, [36]), which recommends selecting the model for which

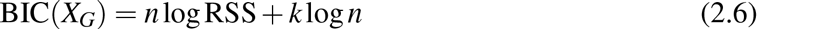

obtains a minimal value. Here RSS is the residual sum of squares under least squares regression, and *k*:= |*G*| refers to the model size. However, in a series of papers ([8, 5, 50]) it was shown that in the context of gene mapping, where *p* is much larger than *n* and the true model is assumed to be relatively small, this criterion leads to a large number of false discoveries. To solve this problem various modifications of the Bayesian Information Criterion were introduced, like mBIC2 ([50], [19])

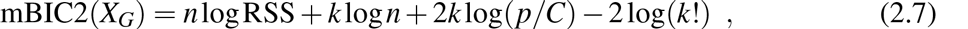

which was designed to control the FDR of wrongly detected SNPs when both *n* and *p* are large but the true number of causal SNPs is relatively moderate. Compared with BIC [36], mBIC2 contains the additional penalty term *2k* log (*p/C*) − 2log(*k*!), which corrects for “multiple testing” and allows to keep the fraction of false discoveries under control in the context of GWAS (see [19]). The criterion is consistent (see for example [43]), thus its FDR converges to zero and the power converges to 1 when the sample size increases. The choice *C* = 4, recommended e.g. in [19], allows to keep FDR below 8% for sample sizes *n* > 200 (see [18] for detailed calculations).

Similarly we will consider linear models for the local ancestry state variables *Z_j_* of the form

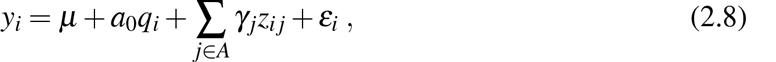

with *A* specifying the set of ancestry markers of the model. We will denote the corresponding design matrix as *Z_A_*. Due to the long range correlation structure of ancestry state variables the corresponding penalty for *Z* variables can be relaxed. A closely related problem occurs for densely spaced markers in experimental crosses discussed in [6], where the selection criterion mBIC was adapted by using an effective number *p^eff^* of markers instead of the total number *p*. Similarly we will here modify mBIC2 for the ancestry state variables

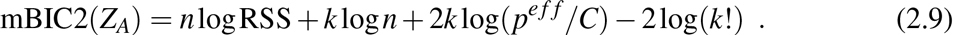

where now*k* = |*A*| is the number of ancestry state variables in the model. The effective number of markers *p^eff^* can be either calculated using a permutation approach or, if the average admixture time is known, based on the theoretical calculations presented in the Appendix.

To fully exploit the potential of admixture mapping a new test statistic was proposed in [46], which combines the genotype and the ancestry information in family based association studies. Here we extend this idea to the case of GWAS in admixed populations, by combining the multiple regression models (2.5) and (2.8) to include both genotypic and ancestry state variables,

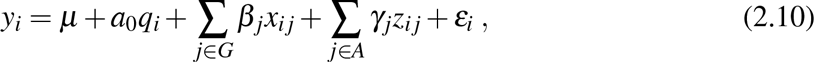

where *G* denotes the set of genotype variables and *A* the set of ancestry state variables included in the model. Accordingly we adapt our model selection criterion to take the form

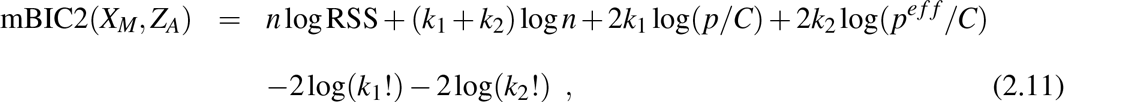

where *k*_1_ is the number of genotype variables included in the model and *k*_2_ is the number of ancestry state variables.

Due to the large number of SNPs considered in GWAS it is not feasible to calculate mBIC2 for all possible regression models. Instead we apply a modification of the classical greedy stepwise approach to search for a model which minimizes the criterion. As a preliminary step we perform single marker tests according to the models (2.3) and (2.4). Step-wise search is then only performed for those markers with p-values smaller than 0.15. Starting from the null model we perform forward selection, where in each consecutive step those markers are added to the model which result in the largest decrease of the mBIC2 criterion. When there are no more markers left which allow to decrease mBIC2 then backward elimination is performed, where in each step that marker is eliminated from the model which gives the largest decrease in mBIC2. Backward elimination is terminated if the removal of any of the remaining markers does not lead to a further decrease of mBIC2. Forward search and backward elimination are performed alternately till convergence.

### 2.2 Simulation Study

To investigate the performance of our model selection strategy we generated data in computer simulations for 1000 individuals from an admixture of the West African (YRI) and European (CEU) populations as discussed in [32]. The ancestry state variables were simulated, based on the hybrid isolation model of [28], for 482 906 autosomal SNPs from the Illumina 650K microarray. The average admixing time was equal to 10 generations, and the average proportion of the genome inherited from YRI was 0.7. The genotype data for the blocks of a given ancestry were obtained by a random selection of individuals of this ancestry from the HapMap Consortium [21] genotype data. Since the average admixture time for the simulated data is known, we can avoid the computationally intensive permutation approach and use instead the theory described in the Appendix to derive the effective number of ancestry markers. The resulting number of *p^eff^* = 4722 tests for ancestry state variables is approximately 100 times smaller than the total number of SNPs.

We consider three different simulation scenarios. In the first scenario trait data were generated under the total null hypothesis according to *y_i_* ~ *N*(0,1). For the two scenarios imitating complex traits we determined 24 SNPs to be “causal” (see (Table 2.1). The 24 selected SNPs all have relatively large minor allele frequencies (*MAF* ≥ 0.4) and can be divided into three groups. Eight of them were strongly correlated with some neighboring SNPs (for each of them the maximum genotypic correlation with fifty neighboring SNPs in each direction exceeded 0.94) and had the same allelic frequencies in both parental populations. The second group consisted of 8 SNPs which were again strongly correlated with neighboring SNPs but now had substantially different allelic frequencies in the two parental populations (difference in frequency of a given allele exceeded 0.7). The last group contained SNPs whose genotypes were only slightly correlated with genotypes of neighboring SNPs while the allelic frequencies in both parental populations were substantially different.

For the second and third scenario trait values for each individual *i* = 1,…, *n* were simulated using the SNPs from Table 2.1 as causal according to the following two multiple regression models:

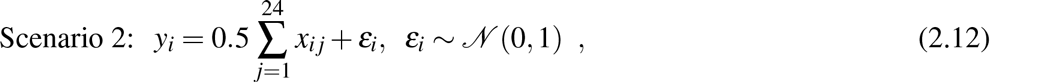

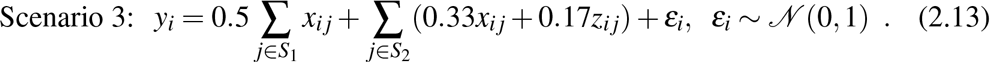

**Table 2.1.**
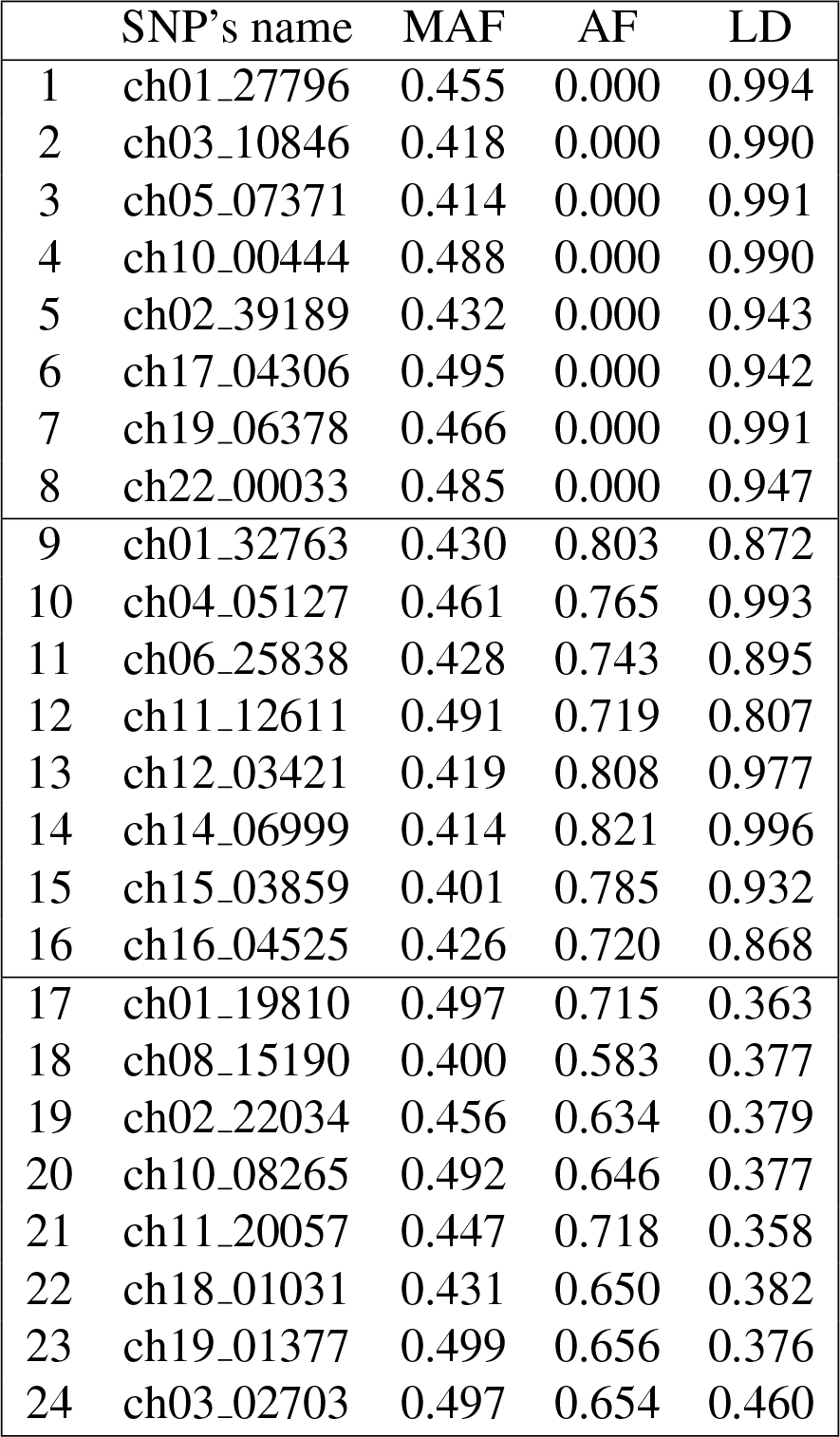
Selected SNPs and their characteristics: MAF is the minor allelic frequency in the admixed population, AF is the difference in allelic frequencies between both admixing populations and LD is defined as the maximum of genotypic correlation with the 100 nearest SNPs (50 on each side).

The sets *S*_1_ and *S*_2_ are defined as *S*_1_ = {1,2,3,4,9,10,11,12,17,18,19,20} and *S*_2_ = {1,…, 24} \ *S*_1_. Simulations according to the last model (2.13) represent the situation when both SNP and ancestry state have an impact on the trait (ie. the SNP efffect is population specific). In each group of 8 we choose half of the SNPs to be population specific. After generating the trait we eliminated the “causal” SNPs from the design matrix to imitate the common situation where the causal mutation has not been genotyped and can only be detected through its neighboring SNPs. The proportion of phenotypic variance explained by all true causal variants is equal to 0.75 in Scenario 2 and 0.73 in Scenario 3.

In our simulation study we investigate the performance of mBIC2 when applied only to genotype variables (2.5), only to ancestry variables (2.8) or when combining both types (2.10). In all cases we use the constant *C* = 4, recommended in [19]. For the classical single-marker tests (2.3) and (2.4) we applied both Bonferroni correction and the more liberal Benjamini Hochberg (BH, [4]) multiple testing procedure. For tests of genotype variables *x_ij_* and of ancestry variables *Z_ij_* we considered Bonferroni adjusted significance levels of 0.05/*p* and 0.05/*p^eff^*, respectively. BH was performed at the same nominal levels, which means that the *k^th^* smallest p-value was compared to 0.05*k*/*p* for genotype variables and to 0.05*k*/*p^eff^* for ancestry state variables, respectively. Finally we compared our approach to *BMIX* [37], which also combines genotype and ancestry information.

In the first experiment we investigated the performance of different methods in the situation when the trait has no genetic component. For this purpose the trait values were independently generated from the standard normal distribution and 1000 replicates were simulated to estimate the Family Wise Error Rate and the average number of false discoveries. In the second and third experiment we simulated 250 independent data sets from the complex models (2.12) and (2.13), respectively, and calculated the average number of true and false discoveries (TP and FP) as well as the empirical false discovery rate (FDR). In case of mBIC2 we define a detected SNP to be a true positive if the correlation between the variable representing this SNP in the identified model and the respective “causal” variable exceeded 0.3. Here we always compare variables of the same type (that is either ancestry state or genotype variables) with each other. When an ancestry variable in the model is strongly correlated with the ancestry variable at the location of the causal SNP we count this detection as a true positive, even when the data generating model included only the genotype variable of that location. Two or more detections corresponding to the same causal SNP are counted as just one true positive, all other detections are classified as false positives.

For the multiple testing procedures one typically observes that detections appear in “clumps” of correlated SNPs, marking the potentially interesting regions of the chromosome. In this case we use the concept of scan statistics to define false and true discoveries [40]. For each of the detected variables (region seed) we form a detection region consisting of other detected variables of the same type whose correlation with the seed exceeds 0.3. When the detection regions of different SNPs intersect, we combine them to form a larger clump. The clump is considered a true discovery if at least one of its members is strongly correlated with the respective causal variable (*ρ* > 0.3), otherwise it is classified as a false discovery.

To analyze the dependency of our procedure on the sample size *n* and the choice of the scaling constant *C* we performed additional simulations under Scenario 2. We consider *n* in the range between 650 and 1000, where smaller samples were obtained by random elimination of individuals from our design matrix. The reported results are based on 250 independent replicates of each experiment.

### 2.3 Analysis of HDL in an African American cohort

We applied the joint genotype-ancestry model to analyze the concentration of high-density lipoprotein (HDL) cholesterol in African Americans, using genotype and phenotype data from the Women’s Health Initiative SNP Health Association Resource (WHI-SHARe). The WHI is a U.S.- based study focusing on common health issues in postmenopausal women. Individual characteristics of the participants and genotyping quality assessment analyses are described in [13]; in total, 656852 SNPs passed all QC criteria. Here we re-analyze the log-transformed HDL phenotype in 8153 individuals. Genome-wide European ancestry proportions of the individuals are estimated using the program frappe [45], while locus specific ancestries are estimated using SABER+ [26]. The number of effective ancestry tests *p^eff^* = 6000 was calculated based on the permutation approach.

## 3 Results

### 3.1 Simulation Study

#### 3.1.1 Scenario 1: Weak sense Family Wise Error Rate

The estimated FWER (probability of detecting at least one signal) based on 1000 replicates for different procedures are presented in Table 3.1. Note that FWER of the Bonferroni procedure for the ancestry single marker tests slightly exceeds the nominal level of 5%. This may be explained by the fact that markers are not uniformly spaced and admixture times vary between different individuals. Both of these distributional effects might not be captured adequately by using the respective average values in our theoretical formulas. However, our model selection approach controls FWER at the desired level. Due to the consistency of mBIC2 and the relatively large sample size the FWER for search over ancestry dummy variables with mBIC2 is as low as 2.3%. Enlarging the design matrix by including genotype state variables leads to an increased FWER of 3.4%, which is still substantially below 5%. The largest FWER is produced by BMIX at approximately 8%. This is still reasonably small given that the “effective number of tests” used by BMIX is equal to 370, which is much smaller than our own estimate of 4722. This is probably because BMIX uses the effective number of tests only in a rather informal manner when constructing a prior probability for the expected number of causal mutations.

**Table 3.1:** Comparison of Family-wise Error Rate (FWER) and expected number of false positives (FP) under the global null hypothesis. Bonf and BH refer to single-marker tests with Bonferroni and Benjamini Hochberg procedure at nominal levels 0.05. mBIC2 refers to the model selection approach. Methods with *X* use only genotypic markers, with *Z* only ancestral markers, and with *X* + *Z* a combination of both types of markers.

|  | Bonf |  | BH |  | BMIX<br>X+Z | mBIC2 |  |  |
| --- | --- | --- | --- | --- | --- | --- | --- | --- |
|  | X | Z | X | Z |  | X | Z | X+Z |
| FWER | 0.044 | 0.069 | 0.044 | 0.069 | 0.080 | 0.010 | 0.023 | 0.034 |
| FP | 0.045 | 0.071 | 0.046 | 0.074 | 0.086 | 0.011 | 0.024 | 0.036 |

**Table 3.2:**
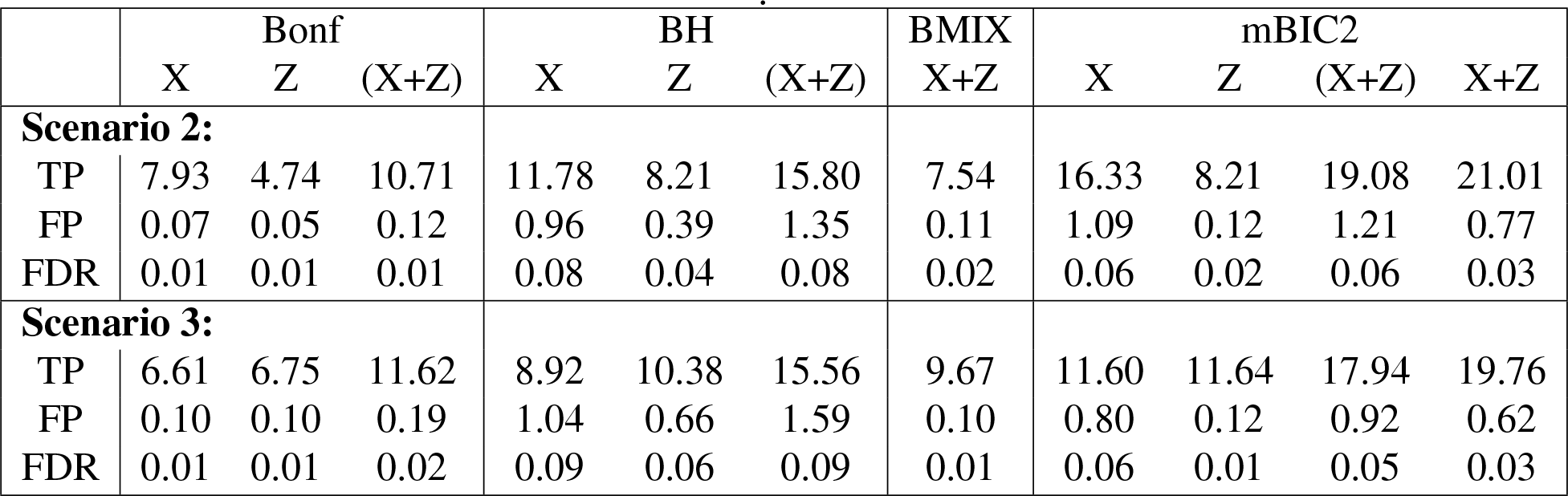
Summary of results for Scenario 2 and 3 in terms of expected true positives (TP), false positives (FP) and false discovery rate (FDR). The columns *(X* + *Z*) present results obtained by combining the separate analysis over genotype and ancestry based GWAS, while the last column *X* + *Z* refers to our new model selection approach based on (2.11)

#### 3.1.2 Scenario 2 and 3: Complex traits

Table 3.2 summarizes the simulation results for the two scenarios with complex traits. We start with looking at the results of the BMIX procedure, which turns out to be in both scenarios more conservative than any of the mBIC2 based selection procedures. This is in contrast to the results reported in Table 3.1 for simulations under the total null hypothesis. For Scenario 3 BMIX has a much smaller Type I error rate than the Bonferroni correction applied to both marker types (*X* + *Z*), which goes along with a substantially decreased power. For Scenario 2 BMIX and Bonferroni correction for (*X* + *Z*) have comparable Type I error rates, but BMIX still has substantially lower power. In contrast mBIC2 for combined genotypic and ancestry state variables has only slightly larger FDR than BMIX, but in Scenario 3 more than twice and in Scenario 2 almost three times larger power.

Concerning single-marker tests, as expected BH has in both scenarios much larger power than the Bonferroni procedure which comes at the price of a substantially larger number of false positives. Comparing mBIC2 with BH for the same type of marker, respectively, one observes that generally speaking mBIC2 has larger or comparable power and at the same time lower FDR. The average number of false positives is for *X* variables at a similar level, whereas for *Z* variables it tends to be smaller for mBIC2. In both scenarios the new mBIC2 procedure which combines genotype and local ancestry information outperforms all other methods in terms of power, while keeping the FDR at a level of 3%.

**Table 3.3:** Power to detect individual SNPs for Scenario 2. Large power is marked by red and small power by blue. In the last column *X* + *Z* the values in brackets give the percentage of cases for which detection resulted from ancestry dummy variables.

|  | Bonf |  | BH |  | BMIX | mBIC2 |  |  |
| --- | --- | --- | --- | --- | --- | --- | --- | --- |
|  | X | Z | X | Z | X+Z | X | Z | X+Z |
| 1 | 0.98 | 0.00 | 1.00 | 0.00 | 0.56 | 1.00 | 0.00 | 1.00 (Z: 0.00) |
| 2 | 0.73 | 0.00 | 0.92 | 0.00 | 0.19 | 0.99 | 0.00 | 1.00 (Z: 0.00) |
| 3 | 1.00 | 0.00 | 1.00 | 0.00 | 0.91 | 1.00 | 0.00 | 1.00 (Z: 0.00) |
| 4 | 0.48 | 0.00 | 0.76 | 0.00 | 0.04 | 0.99 | 0.00 | 0.97 (Z: 0.00) |
| 5 | 1.00 | 0.00 | 1.00 | 0.00 | 0.93 | 1.00 | 0.00 | 1.00 (Z: 0.00) |
| 6 | 0.40 | 0.00 | 0.68 | 0.00 | 0.00 | 1.00 | 0.00 | 1.00 (Z: 0.00) |
| 7 | 0.64 | 0.00 | 0.91 | 0.00 | 0.39 | 1.00 | 0.00 | 1.00 (Z: 0.00) |
| 8 | 0.26 | 0.00 | 0.66 | 0.00 | 0.01 | 0.99 | 0.00 | 1.00 (Z: 0.00) |
| 9 | 0.22 | 0.50 | 0.54 | 0.84 | 0.42 | 0.74 | 0.85 | 0.98 (Z: 0.74) |
| 10 | 0.62 | 0.53 | 0.92 | 0.83 | 0.67 | 0.99 | 0.67 | 1.00 (Z: 0.10) |
| 11 | 0.21 | 0.19 | 0.59 | 0.49 | 0.34 | 0.96 | 0.47 | 0.96 (Z: 0.20) |
| 12 | 0.00 | 0.01 | 0.02 | 0.11 | 0.04 | 0.96 | 0.02 | 0.87 (Z: 0.15) |
| 13 | 0.58 | 0.76 | 0.88 | 0.94 | 0.71 | 1.00 | 0.82 | 1.00 (Z: 0.30) |
| 14 | 0.10 | 0.31 | 0.39 | 0.71 | 0.45 | 0.97 | 0.82 | 1.00 (Z: 0.06) |
| 15 | 0.17 | 0.09 | 0.55 | 0.40 | 0.34 | 0.88 | 0.53 | 0.98 (Z: 0.24) |
| 16 | 0.54 | 0.85 | 0.88 | 0.97 | 0.62 | 1.00 | 0.94 | 1.00 (Z: 0.08) |
| 17 | 0.00 | 0.00 | 0.00 | 0.00 | 0.00 | 0.00 | 0.00 | 0.35 (Z: 0.35) |
| 18 | 0.00 | 0.00 | 0.00 | 0.04 | 0.05 | 0.01 | 0.03 | 0.25 (Z: 0.23) |
| 19 | 0.00 | 0.00 | 0.00 | 0.02 | 0.00 | 0.01 | 0.25 | 0.57 (Z: 0.56) |
| 20 | 0.00 | 0.62 | 0.04 | 0.88 | 0.46 | 0.32 | 0.69 | 0.92 (Z: 0.91) |
| 21 | 0.00 | 0.32 | 0.02 | 0.72 | 0.27 | 0.22 | 0.78 | 0.97 (Z: 0.95) |
| 22 | 0.00 | 0.19 | 0.00 | 0.52 | 0.14 | 0.03 | 0.67 | 0.90 (Z: 0.90) |
| 23 | 0.00 | 0.38 | 0.00 | 0.74 | 0.03 | 0.01 | 0.65 | 0.68 (Z: 0.68) |
| 24 | 0.00 | 0.00 | 0.01 | 0.00 | 0.00 | 0.26 | 0.00 | 0.62 (Z: 0.51) |

We will next discuss the power of different procedures to detect individual SNPs as summarized in Tables 3.3 and 3.4, which provide the percentage of simulation runs for which a corresponding neighborhood was identified. For genotypic markers the selection based on mBIC2 applied separately to *X* has in the majority of cases larger power than BH. An interesting example is provided by SNP number 12 in Scenario 2. This SNP has a relatively large LD (0.807) and is detected with a power of 96% by mBIC2 applied to the genotype data matrix *X*. On the other hand both single-marker tests have very small power to detect this SNP. This phenomenon can be easily explained by the fact that single-marker tests look only at the correlation between the genotype (or the ancestry status) of a given SNP and the trait. However, such correlations do not necessarily represent the strength of the effect of a given SNP but depend also on the correlations between this SNP and other causal variants (see [19] for more details). In fact, the power of detecting a given SNP by single-marker tests depends on the so called non-centrality parameter, which captures these inter-SNP correlations (see Figure 3.1) and is calculated according to the formula

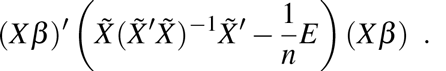

**Table 3.4.** Power to detect individual SNPs for Scenario 3.

|  | Bonf |  | BH |  | BMIX | mBIC2 |  |  |
| --- | --- | --- | --- | --- | --- | --- | --- | --- |
|  | X | Z | X | Z | X+Z | X | Z | X+Z |
| 1 | 1.00 | 0.00 | 1.00 | 0.00 | 0.84 | 1.00 | 0.00 | 1.00 (Z: 0.00) |
| 2 | 0.89 | 0.00 | 0.98 | 0.00 | 0.34 | 0.98 | 0.00 | 0.98 (Z: 0.00) |
| 3 | 1.00 | 0.00 | 1.00 | 0.00 | 0.80 | 1.00 | 0.00 | 1.00 (Z: 0.00) |
| 4 | 0.80 | 0.00 | 0.95 | 0.00 | 0.33 | 0.98 | 0.00 | 0.95 (Z: 0.00) |
| 5 | 0.82 | 0.00 | 0.95 | 0.04 | 0.88 | 0.71 | 0.01 | 0.90 (Z: 0.00) |
| 6 | 0.00 | 0.00 | 0.02 | 0.00 | 0.00 | 0.39 | 0.03 | 0.60 (Z: 0.16) |
| 7 | 0.00 | 0.04 | 0.06 | 0.22 | 0.46 | 0.36 | 0.20 | 0.54 (Z: 0.18) |
| 8 | 0.00 | 0.00 | 0.01 | 0.04 | 0.07 | 0.12 | 0.10 | 0.53 (Z: 0.10) |
| 9 | 0.58 | 0.72 | 0.86 | 0.95 | 0.72 | 0.64 | 0.97 | 0.98 (Z: 0.73) |
| 10 | 0.73 | 0.68 | 0.92 | 0.92 | 0.77 | 0.97 | 0.89 | 1.00 (Z: 0.28) |
| 11 | 0.30 | 0.27 | 0.62 | 0.63 | 0.42 | 0.90 | 0.82 | 0.96 (Z: 0.30) |
| 12 | 0.04 | 0.09 | 0.13 | 0.37 | 0.11 | 0.81 | 0.30 | 0.86 (Z: 0.40) |
| 13 | 0.34 | 0.98 | 0.66 | 1.00 | 0.74 | 0.95 | 0.99 | 1.00 (Z: 0.94) |
| 14 | 0.05 | 0.49 | 0.25 | 0.84 | 0.53 | 0.58 | 0.98 | 1.00 (Z: 0.91) |
| 15 | 0.04 | 0.46 | 0.21 | 0.84 | 0.28 | 0.62 | 0.96 | 0.99 (Z: 0.81) |
| 16 | 0.00 | 0.02 | 0.00 | 0.11 | 0.01 | 0.02 | 0.15 | 0.15 (Z: 0.09) |
| 17 | 0.00 | 0.00 | 0.00 | 0.00 | 0.00 | 0.00 | 0.03 | 0.45 (Z: 0.45) |
| 18 | 0.00 | 0.04 | 0.01 | 0.28 | 0.43 | 0.02 | 0.23 | 0.45 (Z: 0.42) |
| 19 | 0.00 | 0.00 | 0.00 | 0.08 | 0.00 | 0.00 | 0.66 | 0.77 (Z: 0.76) |
| 20 | 0.02 | 0.93 | 0.15 | 0.99 | 0.82 | 0.16 | 0.89 | 0.82 (Z: 0.82) |
| 21 | 0.01 | 0.60 | 0.11 | 0.92 | 0.42 | 0.26 | 0.99 | 0.99 (Z: 0.99) |
| 22 | 0.00 | 0.56 | 0.01 | 0.88 | 0.31 | 0.02 | 0.94 | 0.99 (Z: 0.99) |
| 23 | 0.00 | 0.85 | 0.00 | 0.96 | 0.20 | 0.03 | 0.94 | 0.97 (Z: 0.97) |
| 24 | 0.00 | 0.03 | 0.01 | 0.32 | 0.04 | 0.09 | 0.57 | 0.88 (Z: 0.87) |

Here *X* denotes the matrix containing genotypes of all causal SNPs,*β* is the vector of true regression coefficients, 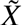 contains all variables included in the single-marker test model (2.3) (i.e. the columns of ones, *qi* and *x_ij_*), where the causal mutation is replaced by the most correlated SNP which was genotyped and *E_n×n_* is the matrix for which all elements are equal to 1.

**Figure 3.1:**
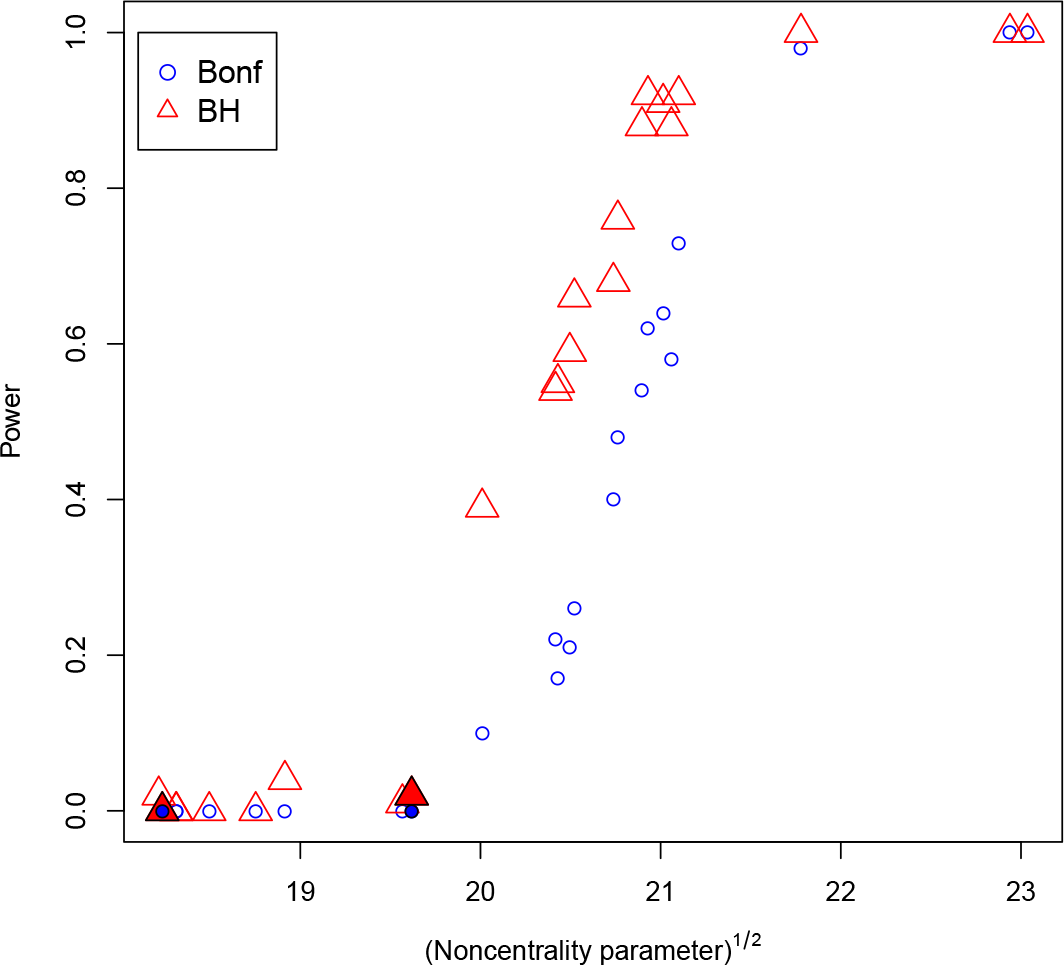
Power for individual tests (matrix X) vs. noncentrality parameter for Scenario 2. Filled red triangles and filled blue circles mark two SNPs discussed in the text: SNP12 (low power for Bonf and BH but high power for mBIC2) is near the middle of the picture,SNP17 (high AF but seen only for X+Z) on the left.

Figure 3.1 shows that for SNP number 12 the non-centrality parameter takes a rather small value, and thus it cannot be detected by single-marker tests. Instead, it is rather easily detected within the framework of multiple regression, which takes into account the effects of other causal SNPs. In Figure 3.1 we also highlight SNP number 17, which has high AF (0.715), but when we use only the matrix Z, even mBIC2 has no power to detect it (the non-centrality parameter for this SNP is the lowest of all). However, it was found in 35% of the simulation runs when both matrices are used.

Looking at the results specifically in terms of the three different types of causal SNPs used in the simulation study, then Table 3.3 indicates that combining genotype and ancestry data increases power particularly for mutations in regions of low LD. In the combined analysis of model (2.10) SNPs of the group 17 - 24 were almost exclusively detected by ancestry state variables, and in almost all cases the power to detect these SNPs was substantially lower when using model (2.8) with the *Z* matrix alone. The explanation for this is that in the combined model the effect of detected genotype variables is removed from the residual error and thus it becomes easier to detect further ancestry state variables. For Scenario 3 the gain in power for the last 8 SNPs by combining genotype and ancestry data is smaller somewhat than in Scenario 2 but there is additional gain in power obtained for SNPs 5 - 8. These SNPs of the first group are not easily detected when searching over *X* or *Z* variables separately, whereas the combined approach yields substantial power for all four SNPs (especially for SNP 8). The reason for this gain in power is the very same as in Scenario 2. By including many other causal SNPs in the model the residual sum of squares is reduced and the chance is increased to detect these SNPs. Comparing Scenario 2 with Scenario 3 furthermore shows that ancestry variables play a substantial role in identifying mutations whose effect is population specific. In Scenario 2 SNPs 13-16 are mainly detected by *X* variables, whereas in Scenario 3 they are identified mainly with *Z* variables.

The final example of this section will discuss the working principle of our new approach to combine genotypic and ancestry state data. Figure 3.2 illustrates the role of *X* and *Z* in detecting SNP 14 in Scenario 2 as a function of the sample size. For smaller sample sizes this SNP is detected mainly by the ancestry state variable *Z*, whereas for larger *n* it is more frequently identified by the genotype variable *X*. This is due to the fact that for small sample sizes the reduced multiple testing correction used in admixture mapping substantially enhances the power. Instead, when the sample size is large enough, the causal gene can be detected by the genotype variable, which provides a more precise localization. Note that in this argument the sample size can be substituted by the magnitude of the gene effect. Consequently in our admixture mapping approach the ancestry state variables are helpful for detecting genes with small effects, which however comes at the price of rather imprecise estimation of their location. The major advantage of our combined mBIC2 criterion (2.11) is that it allows to decide automatically for every SNP whether selection is based on *X* or *Z* variables, depending on the magnitude of its effect and on sample size.

**Figure 3.2:**
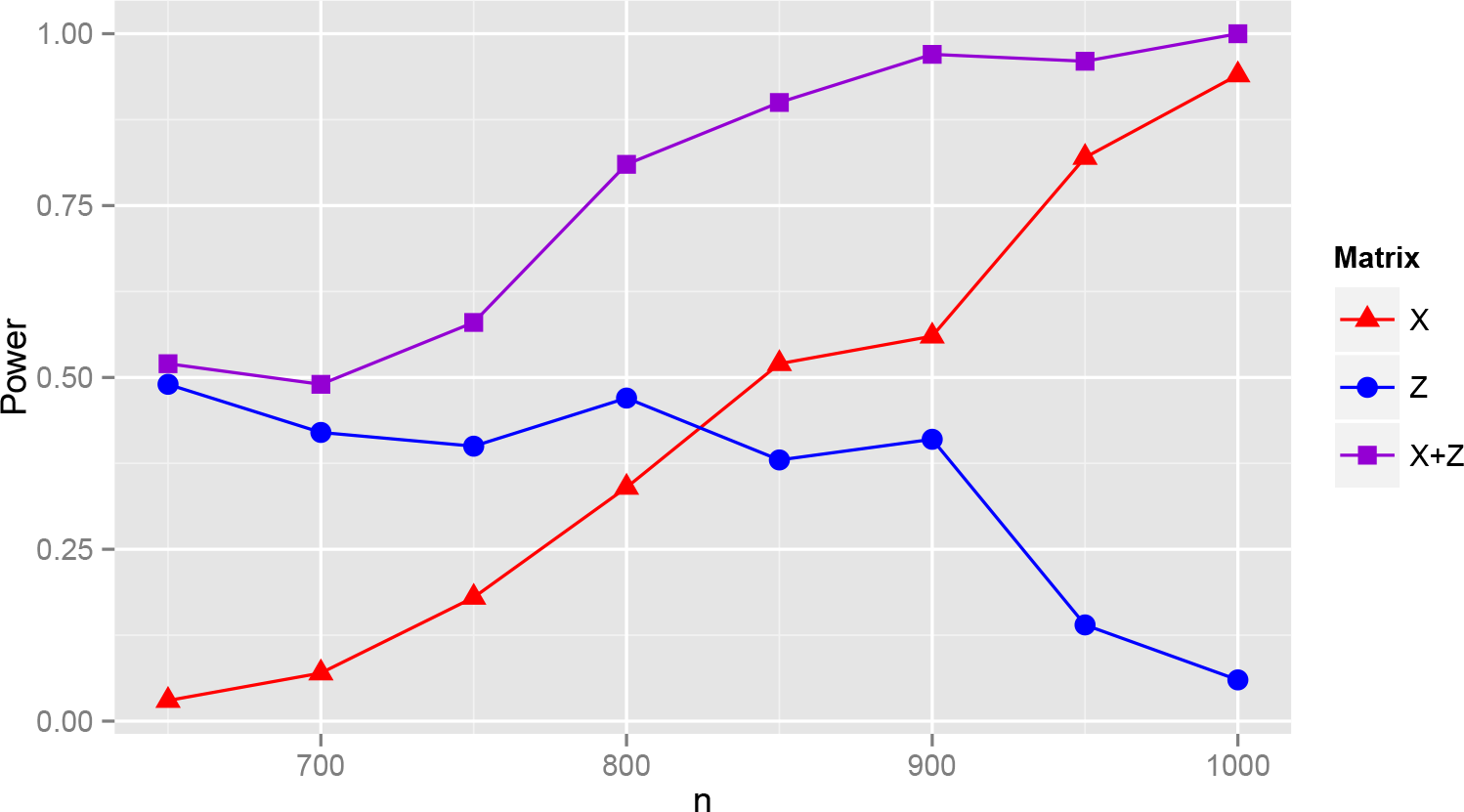
Power for SNP 14 in Scenario 2 for different sample size n based on model selection with mBIC2 using both genetic and ancestry markers (*X* + *Z*). Here the lines denoted by *X* and *Z* illustrate the proportion of detections due to genetic markers and ancestry markers, respectively, for the combined approach.

### 3.2 WHI-SHARe HDL analysis

The joint genotype-ancestry analysis produces a multivariate model that includes seven genotype variables and two locus-specific ancestry variables. Table 3.5 lists these variables in the order they enter the model, as well as the single-marker p-values from either genotype-based or admixture mapping analysis. The first seven terms to enter the multivariate model, including six SNP genotype variables and the locus-specific ancestry on chromosome 11, coincided with the genome-wide significant findings through single-marker analyses [13]. Interestingly, the next term to enter the multivariate model was the local ancestry on chromosome 17q, which does not meet genome-wide significance in a single-marker admixture mapping analysis (*p* = 5.80 ~ 10^−5^). In contrast, the local ancestry at 9q22, which is significant in admixture map-ping (*p* = 5.58 × 10^−7^) was not selected by the multivariate model. We verified that given the first seven variables entered in the model, adding local ancestry on 17q indeed led to a greater improvement in model fitting than adding the local ancestry of 9q22: the multiple R-squared statistics were 0.04884 and 0.04854, respectively. Of course, without an independent validation data, we cannot say which model will have better predictive value; however, these results illustrate that a multivariate model can prioritize variables according to a different order than a single-variable approach. Finally, the last variable to enter the model, SNP rs7249565 has a p-value of 1.13 × 10^−5^ in a single-marker test; after including all the other eight variables, its p-value (corresponding to the main effect of this SNP in the multivariate model) decreased to 1.10*e* − 07, presumably because the other variables reduced the estimated residual variance.

**Table 3.5:** Results of real data analysis

|  | Single SNP p-value | Single AM p-value | p-value in the multivariate model |
| --- | --- | --- | --- |
| CH16rs247617 | 1.48E-44 | | $< 2e-16$ |
| CH7rs6963015 | 7.53E-10 |  | 7.96E-11 |
| CH8rs326 | 1.23E-08 |  | 8.92E-09 |
| CH8rs1461729 | 7.39E-09 |  | 1.04E-08 |
| CH21rs13046373 | 2.26E-08 |  | 3.35E-08 |
| CH19rs12979813 | 1.99E-09 |  | 1.79E-09 |
| CH11rs531964anc |  | 2.82E-07 | 2.05E-06 |
| CH17rs11867417anc |  | 5.80E-05 | 4.38E-06 |
| CH19rs7249565 | 1.13E-05 |  | 1.10E-07 |

## 4 Discussion

The presented simulation study and real data analysis demonstrate that our model selection approach allows for efficient integration of genetic and ancestry information. As expected, in comparison to classical GWAS inclusion of ancestry state variables results in a substantial increase of power to detect causal mutations in the regions of low LD. Also, in comparison to admixture mapping, our approach allows for substantially larger power to detect “admixture effects”, due to reducing the residual error by including detected genotype state variables in the model. Interestingly, inclusion of admixture state variables usually does not deflate the power of detecting genes in regions of high LD, because the long range correlations between admixture variables make it possible to use a relatively small additional correction for multiple testing. The corresponding increase in the penalty of the selection criterion is most often counterbalanced by the reduction of the residual error due to detected admixture variables. Furthermore our simulation study shows that the admixture state variables help to detect causal mutations in regions of high LD, if the gene signal is weak or the sample size is small. The presented model selection approach provides an automatic choice between genotype and ancestry state variables, which yields high power of gene detection if the sample size is small (choice of admixture variable) and high precision of gene localization if the sample size is large enough (choice of genotype variable).

Our model selection approach creates a general framework for GWAS in admixture populations, which, apart from other advantages, allows to incorporate the dependency of gene effects on the population specific genetic background. The search procedure gives a high power of gene detection while keeping the number of false discoveries under control. The approach can be easily extended for case-control studies and any other trait distribution which can be modelled by Generalized Linear Models (see[50] or [14]). According to [49] one can also expect good performance of a rank based version of our criterion in case when the trait distribution is heavy tailed or the data contains some proportion of outliers. These assertions will be investigated in some follow-up research. R code which has implemented the presented procedure for GWAS in admixed population is available at http://prac.im.pwr.wroc.pl/~mbogdan/admixtures/ and will soon be expanded to include the analysis of case-control studies, GLM and a rank based version.

## Acknowledgements

PS and MB were supported by the European Union’s 7th Framework Programme for research, technological development and demonstration under Grant Agreement no 602552, co-financed by the Polish Ministry of Science and Higher Education under Grant Agreement 2932/7.PR/2013/2. HT was supported by the US National Institutes of Health grants GM073059.

## A Calculation of the effective number of ancestry tests in the simulation study

Firstly, assume that we perform a genome scan based on ancestry markers which are equally spaced at a distance of *L* Morgan. To assess the necessary multiple testing correction for the ancestry markers we consider the simple multiple regression models (2.4) where we are simultaneously testing the null hypotheses *H_0j_*: *γ_j_* =0. Consider an individual with admixing time *t* and a genome-wide ancestry value *q_i_*. Elementary calculations show that the conditional correlation between ancestry state variables at the neighboring loci does not depend on the specific ancestry value, but only on the individual admixing time and is given by

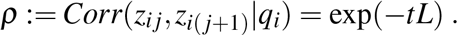

Moreover, based on the arguments presented in [17] (see also [39]), the sequence of test statistics at consecutive locations can be approximated by the square of an Ornstein-Uhlenbeck process. One then can show that the family wise error rate *a* of such a search is approximately

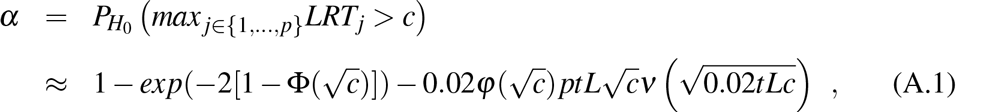

where Φ(·) and φ(·) denote the cumulative distribution function and the density of the standard normal distribution and

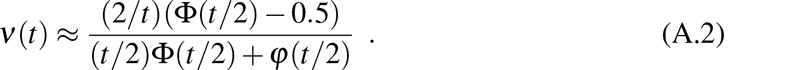

On the other hand, the family wise error rate resulting from performing *p^eff^* independent tests is equal to

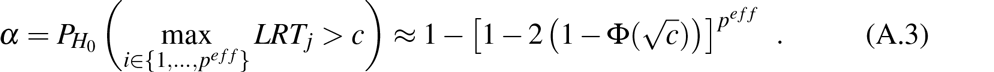

Comparing (A.1) and (A.3) results in the following effective number of tests for the ances try state variables

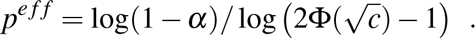

As observed in [6] the dependency of *p^eff^* on *α* is rather weak and the value of *p^eff^* calculated for *α* = 0.05 can be used as a good approximation for *p^eff^* corresponding to any *α* ∈ (0,0.1]. In the simulation study we calculated the effective number of tests separately for each chromosome. Since the average admixture time is equal to 10, for this analysis we replaced *tL* in equation (A.1) with 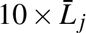, where 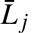 is the average distance between neighboring markers on a given chromosome. Finally,the effective number of SNPs for mBIC2 was obtained by adding the effective number of tests on each chromosome, which results in *p^eff^* = 722.

## References

[1] Balding D.J. (2006) A tutorial on statistical methods for population association studies Nat. Rev. Gen. 7:781–791.

[2] Baierl A., Bogdan M., Frommlet F., Futschik A. (2006) On locating multiple interacting Quantitative Trait Loci in intercross designs. Genetics 173: 1693–1703.

[3] Baierl A., Futschik A., Bogdan M., Biecek P. (2007) Locating multiple interacting quantitative trait loci using robust model selection. Computational Statistics and Data Analysis 51: 6423–6434.

[4] Benjamini, Y. and Hochberg, Y. (1995). Controlling the false discovery rate: a practical and powerful approach to multiple testing. J. Roy. Statist. Soc. Ser. B. 57, 289–300. MR1325392

[5] Bogdan M., Ghosh J.K., Doerge, R.W. (2004). Modifying the Schwarz Bayesian Information Criterion to locate multiple interacting quantitative trait loci. Genetics 167: 989–999.

[6] Bogdan M., Frommlet F., Biecek P., Cheng R., Ghosh J.K., Doerge R.W. (2008). Extending the Modified Bayesian Information Criterion (mBIC) to dense markers and multiple interval mapping. Biometrics 64: 1162–1169.

[7] Bogdan, M., Zak-Szatkowska, M., Ghosh, J.K. Selecting explanatory variables with the modified version of Bayesian Information Criterion, Quality and Reliability Engineering International, 24: 627–641, 2008.

[8] Broman, K.W., Speed T.P. (2002) A model selection approach for the identification of quantitative trait loci in experimental crosses. J Roy Statist Soc Ser B 64(4): 641656.

[9] Cardon L.R., Palmer L.J. (2003) Population stratification and spurious allelic association. Lancet 361: 598–604.

[10] Chen H.S., Zhu X., Zhao H., Zhang S. (2003) Qualitative semi-parametric test for genetic associations in case-control designs under structured populations. Ann Hum Genet. 67: 250–264.

[11] Chen, J., Chen Z. (2008). Extended Bayesian Information criteria for model selection with large model spaces. Biometrika 95(3), 759–771.

[12] Clarke G., Whittemore A.S. (2007) Comparison of admixture and association mapping in admixed families. Genet Epidemiol 31:763–775.

[13] Coram M.A., Duan Q., Hoffmann T.J., Thornton T., Knowles J.W., Johnson N.A., Ochs-Balcom H.M., Donlon T.A., Martin L.W., Eaton C.B., Robinson J.G., Risch N.J., Zhu X., Kooperberg C., Li Y., Reiner A.P., Tang H. (2013) Genome-wide characterization of shared and distinct genetic components that influence blood lipid levels in ethnically diverse human populations. Am J Hum Genet. 92(6):904–16.

[14] Dolejsi E., Bodenstorfer B., Frommlet F. (2014) Analyzing genome-wide association studies with an FDR controlling modification of the Bayesian information criterion. PLOS ONE 9(7): e103322.

[15] Dupuis J., Siegmund D. O. (1999). Statistical methods for mapping quantitative trait loci from a dense set of markers. Genetics 151: 373–386.

[16] Erhardt, V., M. Bogdan and C. Czado (2010). Locating multiple interacting quantitative trait loci with the zero-inflated generalized Poisson regression, Statistical Applications in Genetics and Molecular Biology, Vol 9: Iss. 1, Article 26.

[17] Feingold, E., P.O. Brown and D. Siegmund: Gaussian models for genetic linkage analysis using complete high resolution maps of identity-by-descent. Am. J. Hum. Genet. 53: 23451 (1993).

[18] Frommlet, F., Bogdan, M. and Chakrabarti, A. (2010). Asymptotic Bayes optimality under sparsity of selection rules for general priors. Technical report, arXiv:1005.4753.

[19] Frommlet F., Ruhaltinger F., Twarog P., Bogdan M. (2012) A model selection approach to genome wide association studies, Computational Statistics and Data Analysis, 56: 1038–1051.

[20] Halder I., Shriver M. (2003) Measuring and using admixture to study the genetics of complex diseases. Hum Genet 1:52–62.

[21] The International HapMap Consortium. (2007). A second generation human haplotype map of over 3.1 million SNPs. Nature 449, 85162.

[22] Hirschhorn J.N., Daly M.J. (2005) Genome-wide association studies for common diseases and complex traits. Nat Rev Genet. 6:95–108.

[23] Hoffman G.E., Logsdon B.A., Mezey J.G. (2013) PUMA: a unified framework for penalized multiple regression analysis of GWAS data. Plos Comput Biol 9(6): e1003101.

[24] Hoggart, C. J., Shriver M. D., Kittles R. A., Clayton D. G., McKeigue P. M. (2004) Design and analysis of admixture mapping studies. Am J Hum Genet. 274 (5): 965–78.

[25] Hoggart C.J., Whittaker J.C., De Iorio M., Balding D.J. (2008) Simultaneous Analysis of All SNPs in Genome-Wide and Re-Sequencing Association Studies. Plos Genet 4(7): e1000130.

[26] Johnson N.A., Coram M.A., Shriver M.D., Romieu I., Barsh G.S., London S.J., Tang H. (2011) Ancestral components of admixed genomes in a Mexican cohort. PLoS Genet. 7(12):e1002410.

[27] Kang H.M., Sul J.H., Service S.K., Zaitlen N.A., Kong S.Y., Freimer N.B., Sabatti C., Eskin E. (2010) Variance component model to account for sample structure in genome-wide association studies. Nat. Genet. 42:348–54

[28] Long J.C. (1991). The genetic structure of admixed populations. Genetics 127, 417–28.

[29] McCarthy M.I., Abecasis G.R., Cardon L.R., Goldstein D.B., Little J., Ioannidis J.P., Hirschhorn J.N. (2008) Genome-wide association studies for complex traits: consensus, uncertainty and challenges. Nat Rev Genet. 9:356–69.

[30] Price A.L., Patterson N.J., Plenge R.M., Weinblatt M.E., Shadick N.A., Reich D. (2006) Principal components analysis corrects for stratification in genome-wide association studies. Nat Genet. 38(8):904–9.

[31] Price A.L., Patterson N., Yu F., Cox D.R., Waliszewska A., McDonald G.J., Tandon A., Schirmer C., Neubauer J., Bedoya G., Duque C., Villegas A., Bortolini M.C., Salzano F.M., Gallo C., Mazzotti G., Tello-Ruiz M., Riba L., Aguilar-Salinas C.A., Canizales-Quinteros S., Menjivar M., Klitz W., Henderson B., Haiman C.A., Winkler C., Tusie-Luna T., Ruiz-Linares A., Reich D. (2007). A genomewide admixture map for Latino populations. Am. J. Hum. Genet. 80:1024–1036.

[32] Price A. L., Patterson N. J., Hancks D.C., Myers S., Reich D., Cheung V. G. and Spielman R. S. (2008). Effects of cis and trans genetic ancestry on gene expression in African Americans. PLOS Genet. 4(12), e1000294.

[33] Price A.L., Tandon A., Patterson N., Barnes K.C., Rafaels N., Ruczinski I., Beaty T.H., Mathias R., Reich D., Myers S. (2009) Sensitive detection of chromosomal segments of distinct ancestry in admixed populations. PLoS Genet 5:e1000519.

[34] Redden D.T., Divers J., Vaughan L.K., Tiwari H.K., Beasley T.M., Fernandez J.R., Kimberly R.P., Feng R., Padilla M.A., Liu N., Miller M.B., Allison D.B. (2006) Regional admixture mapping and structured association testing: conceptual unification and an extensible general linear model. PLoS Genet 2:e137.

[35] Sankararaman S., Kimmel G., Halperin E., Jordan M.I. (2008) On the inference of ancestries in admixed populations. Genome Res 18:668–675.

[36] Schwarz, G. (1978). Estimating the Dimension of a Model. Ann. Statist. 6(2), 461–464.

[37] Shriner D., Adeyemo A., Rotimi C.N. (2011) Joint Ancestry and Association Testing in Admixed Individuals. PLoS Comput Biol 7(12): e1002325. doi:10.1371/journal.pcbi.1002325

[38] Siegmund D. (2004) Model selection in irregular problems: Applications to mapping quantitative trait loci. Biometrika 91: 785–800.

[39] Siegmund D., Yakir B.: The statistics of Gene Mapping, Springer Series in Statistics for Biology and Health, Springer 2007.

[40] Siegmund D., Yakir B., Zhang, N.R. (2011) The false discovery rate for scan statistics. Biometrika 98 (4): 979–985.

[41] Smith M.W., Patterson N., Leutenberger J.A., Truelove A.L., McDonald G.J., Waliszewska A., Kessing B.D., Malasky M.J., Scafe C., Le E., De Jager P.L., Mignault A.A., Yi Z., De The G., Essex M., Sankale J.L., Moore J.H., Poku K., Phair J.P., Goedert J.J., Vlahov D., Williams S.M., Tishkoff S.A., Winkler C.A., De La Vega F.M., Woodage T., Sninsky J.J., Hafler D.A., Altshuler D., Gilbert D.A., O’Brien S.J., Reich D. (2004) A high-density admixture map for disease gene discovery in African Americans. Am J Hum Genet. 74: 1001–1013.

[42] Sundquist A., Fratkin E., Do C.B., Batzoglou S. (2008) Effect of genetic divergence in identifying ancestral origin using HAPAA. Genome Res 18:676–682.

[43] Szulc P. (2012) Weak consistency of modified versions of Bayesian Information Criterion in a sparse linear regression. Probability and Mathematical Statistics 32:47–55.

[44] Tang H., Coram M., Wang P., Zhu X., Risch N.J. (2006) Reconstructing genetic ancestry blocks in admixed populations. Am J Hum Genet 79:1–12.

[45] Tang H., Peng J., Wang P., Risch N.J. (2005) Estimation of individual admixture: analytical and study design considerations. Genet Epidemiol. 28(4):289–301.

[46] Tang H., Siegmund D.O., Johnson N.A., Romieu I., London S.J. (2010) Joint testing of genotype and ancestry association in admixed families. Genet Epidemiol 34: 783–791.

[47] Tian C., Hinds D.A., Shigeta R., Kittles R., Ballinger D.G., Seldin M.F. (2006) A genomewide single-nucleotide-polymorphism panel with high ancestry information for African American admixture mapping. Am J Hum Genet 79:640–649.

[48] Winkler,C.A., Nelson, G.W., Smith, M.W. (2010) Admixture Mapping Comes of Age. Annu. Rev. Genomics Hum. Genet. 11: 65–89

[49] Zak M., Baierl, A., Bogdan M., Futschik A. (2007) Locating multiple interacting quantitative trait loci using rank-based model selection, Genetics, 176: 1845–1854.

[50] Zak-Szatkowska M., Bogdan M. (2011) Modified versions of Bayesian Information Criterion for sparse Generalized Linear Models, Computational Statistics and Data Analysis, 55: 2908–2924.

[51] Zhu, X., Tang, H., Risch, N. (2008) Admixture Mapping and the Role of Population Structure for Localizing Disease Genes. Advances in Genetics 60: 547–569

